# Charge screening and hydrophobicity drive progressive assembly and liquid-liquid phase separation of reflectin protein

**DOI:** 10.1101/2024.11.26.625517

**Authors:** Reid Gordon, Robert Levenson, Brandon Malady, Yahya Al Sabeh, Alan Nguyen, Daniel E. Morse

## Abstract

The intrinsically disordered reflectin proteins drive tunable reflectivity for dynamic camouflage and communication in the recently evolved *Loliginidae* family of squid. Previous work revealed that reflectin A1 forms discrete assemblies whose size is precisely predicted by protein net charge density (NCD) and charge screening by the local anion concentration. Using dynamic light scattering (DLS), Forster resonant energy transfer (FRET) and confocal microscopy, we show that these assemblies, of which 95-99% of bulk protein in solution is partitioned into, are dynamic intermediates to liquid protein-dense condensates formed by liquid-liquid phase separation (LLPS). Increasing salt concentration drives this progression by anionic screening of the cationic protein’s Coulombic repulsion, and by increasing the hydrophobic effect which tips the balance between short-range attraction and long-range repulsion (SALR) to drive protein assembly and ultimately LLPS. Measuring fluorescence recovery after photobleaching (FRAP) and droplet fusion dynamics, we demonstrate that reflectin diffusivity in condensates is tuned by salt concentration and protein NCD. These results illuminate the physical processes governing reflectin A1 assembly and LLPS, and demonstrate the potential for reflectin A1 condensate-based tunable biomaterials. They also compliment previous observations of liquid phase separation in the Bragg lamellae of activated iridocytes and suggest that LLPS behavior may serve a critical role in governing the tunable and reversible dehydration of the membrane-bounded Bragg lamellae and vesicles containing reflectin in biophotonically active cells.

## Introduction

The skin of cephalopods contains a complex optical system combining both pigmentary and structural coloration working in tandem (1–3). Iridocytes and leucophore cells produce structural color using sub-wavelength nanostructures (4). Unlike broad-band scattering leucophores, iridocytes reflect light in an angle- and wavelength-dependent manner via extensive and highly regular membrane invaginations, forming stacks of Bragg lamellae densely filled with the reflectin proteins (5–8). Uniquely, the recently evolved *Loliginidae* squid possess reversibly and finely tunable leucophores and iridocytes. In the iridocytes, neuronally released acetylcholine binds muscarinic receptors, activating a signal transduction pathway (9) that results in the phosphorylation of reflectin proteins, neutralizing Coulombic repulsion of these cationic, intrinsically disordered proteins to drive their resulting folding and hierarchical assembly (5, 6, 10). It has been hypothesized that this assembly drives the cells’ colligative, osmotic dehydration (11), an effect augmented by steric dehydration and by the release of counterions from the folding and condensation of reflectins resulting in Gibbs-Donnan equilibration, all combining to drive a net efflux of water out from the Bragg lamellae upon activation and a net influx upon reversal (12). This reversible dehydration proportionally increases the refractive index of the Bragg lamellae and reduces their thickness and spacing, thus progressively tuning the wavelength and increasing the intensity of reflected light (5, 7, 8, 11, 12). Essentially the same mechanism controls the tunable dehydration and broad-band reflectivity of the reflectin-containing vesicles in the tunably white leucophores (13, 14).

Reflectin proteins are enriched in arginine, aromatic residues and methionine while being devoid of lysine and aliphatic residues (6, 10, 15, 16). They are block-copolymers of cationic linkers and the strongly conserved reflectin repeat motifs (RMs) and N-terminal sequence (RMn) (16, 17); these motifs are methionine-rich and show little sequence similarity to non-reflectin proteins **(Figure S1)** (16, 17). The abundance of RY and GRY sequences in reflectin A1 from *Doryteuthis opalesc*ens suggests that cation-pi interactions provide a major driving force in reflectin folding and assembly (18, 19). Similarly, the enrichment of methionine residues suggests that sulfur-pi bonds could contribute to inter- and intra-protein interactions (18, 20), which is supported by solution NMR spectroscopy of a folded, truncated reflectin peptide (20). Increasing pH progressively deprotonates the abundant histidine residues in reflectin A1, reducing the number of positive charges and acting as an *in vitro* surrogate for the charge neutralization resulting from phosphorylation (11, 17). This reduction in protein net charge density (NCD) causes the reversible folding, as characterized by the formation of CD signatures of secondary structure and disappearance of the disordered signature, and proportional assembly of reflectin A1 into higher order multimers. (11, 17). Altering reflectin A1 protein net charge density by pH titration of histidine residues, genetic insertion of charged residues, and electrochemical titration reveals that the NCD of the cationic linker regions opposes assembly, and progressively reducing their Coulombic repulsion increases assembly size (11, 21, 22). Reducing protein NCD by genetic insertion of negatively charged glutamate residues, electrochemical titration of histidine residues, as well as ionic screening of Coulombic repulsion all similarly drive proportional reflectin A1 assembly (17, 21–23). This remarkable *in vivo* and *in vitro* tunability of reflectin behavior has drawn much bioengineering interest for creating optically tunable biomaterials (24–29).

Reflectin A1 exhibits several features consistent with proteins that undergo liquid-liquid phase separation (LLPS) (18, 19, 30, 31); it is an intrinsically disordered protein, rich in arginine and tyrosine, and its sequence exhibits a pronounced regularity of charge patterning. LLPS of proteins into protein-dense and protein-dilute phases has been demonstrated to form membraneless organelles in living cells that often are regulated by post-translational modifications such as phosphorylation (32–35). These liquid-like condensates can act to sequester other proteins and RNA, organize and regulate signaling complexes, and buffer intercellular protein concentrations (36–38). Acetylcholine-activated reflectin phosphorylation drives reflectin LLPS *in vivo*, demonstrated by ultrastructural changes in the Bragg lamellae and leucocyte vesicles where discrete 10-50 nm diameter particles transition to a uniform, densely staining liquid (5, 13, 39). Isolated Bragg lamellae from activated iridocytes of the Loliginid squid *Lolliguncula brevis* demonstrated liquid properties such as deformability and surface wetting (39). Additionally, recent simulations based on short-angle x-ray scattering (SAXS) of reflectin A1 demonstrate a transition from oligomeric assemblies to LLPS (40).

Here, we demonstrate using Forster resonance energy transfer (FRET) and dynamic light scattering (DLS) that reflectin A1 assemblies are dynamic, and driven by reduction in the protein’s net charge density, as previously demonstrated using other methods (11, 23). Further, using confocal microscopy and fluorescence recovery after photobleaching (FRAP), we reveal that increasing salt concentration drives reflectin A1 LLPS to form protein-dense liquid condensates whose material properties are tuned by protein net charge density. We show that as a function of salt concentration, reflectin assembly is intermediate to LLPS and propose that assemblies and liquid condensates represent stages in a continuum of the same physical process mediated by short-range attraction and long-range repulsion (SALR). We also demonstrate that reflectin A1 phase transitions differ markedly from the recently described process of percolation-coupled phase separation (41). We explore the dynamics of reflectin A1 assemblies and the driving forces of reflectin A1 phase transitions as a biophysical model for Bragg lamellar condensation, which governs their tunable photonic behavior. We investigate the usefulness of reflectin A1 condensates as a basis for tunable biomaterials by probing their time- and net charge density-dependent material properties.

## Results

### Reflectin A1 assemblies coexist with dilute phase

Following assembly of reflectin A1, a centrifugation assay was used to demonstrate and analyze the coexistence of a dense phase of assemblies and a dilute phase of monomers or small oligomers (41). Reflectin A1 at an initial concentration of 100 µM in acetic acid buffer (pH 4, 25 mM) was assembled by dilution to into MOPS buffer (pH 7, 25 mM), and after 5 d at 20°C, samples were centrifuged at 20,000 x g for 6 hours to remove assemblies **(Figure 1A)**. The assemblies were incubated for 5 days to ensure that the two phases had reached equilibrium (42). Bradford assays revealed that the protein concentration in the dilute phase, C*_dilute_,* increased in linear proportion to the bulk protein concentration, C*_bulk_,* for C*_bulk_* between 1-10 µM **(Figure 1B).** Although the protein concentration of the dense assembly phase, C*_dense_*, is unknown, the ratio of moles protein in monomer or small oligomers to moles protein in assemblies was constant for C*_bulk_* ≥ 2 μM. This value decreased from 99.4 ± 2.7% for C*_bulk_*= 1 μM to 94.7 ± 0.1% for C*_bulk_*= 2 μM and remained relatively constant up to C*_bulk_*= 10 μM (95.4 ± 0.18%) **(Figure 1C)**. C_dilute_ for reflectin A1 assemblies depends on C_bulk_, which differs from classical LLPS in which C*_dilute_* increases with C*_bulk_* until it exceeds the saturation concentration C*_sat_*, after which any additional increase in C*_bulk_* is partitioned into the dense phase, after which C*_sat_* = C*_dilute_* (19, 43).

**Figure 1.**
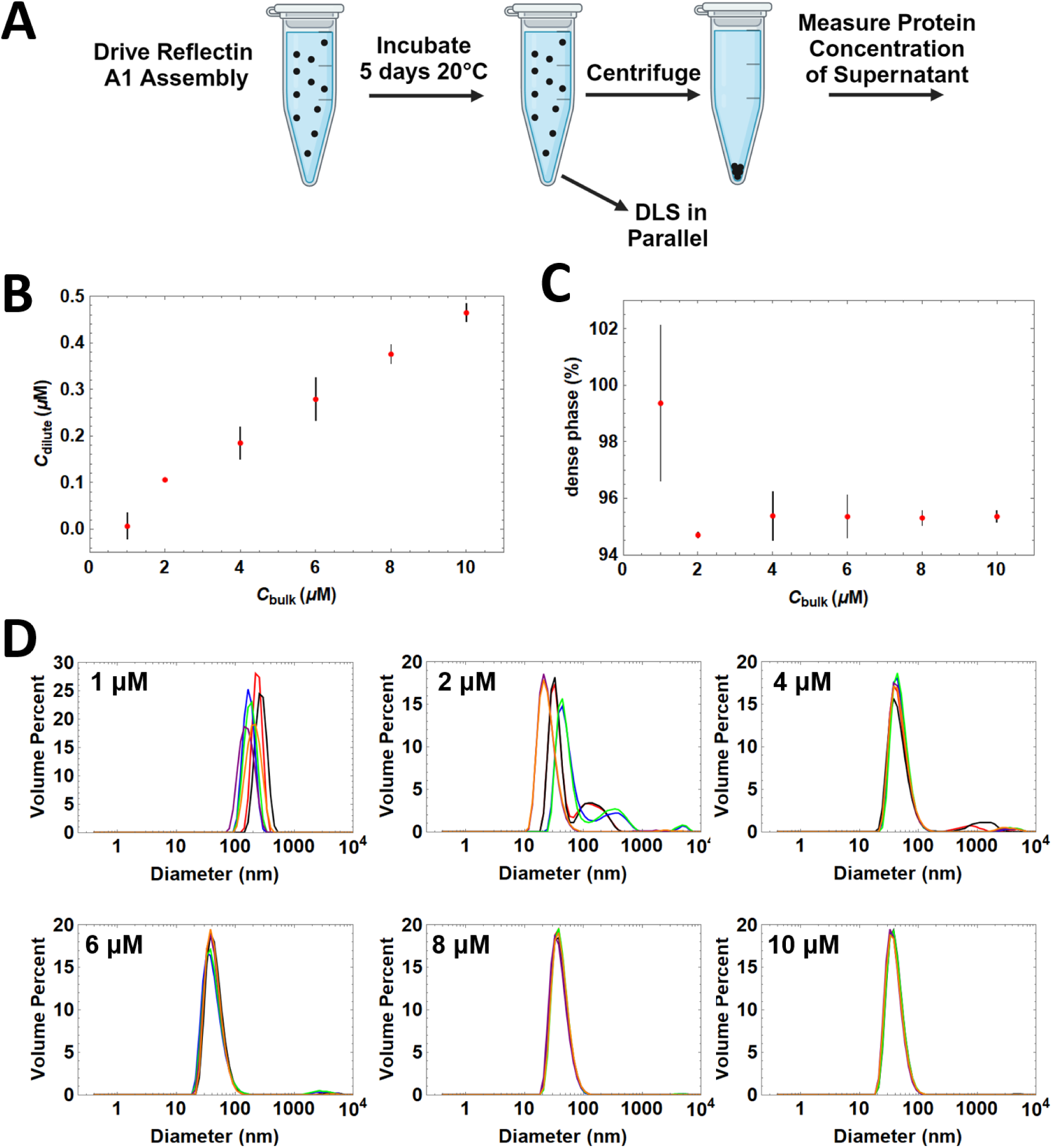
Bradford Assay and DLS of reflectin A1 assemblies as a function of concentration. A) Reflectin A1 was driven to assemble by dilution into MOPS buffer (pH 7 25mM), and after 5 days dilute phase protein concentration determined by centrifugation assay. DLS of assemblies performed in parallel. B) Reflectin A1 dilute concentration (y-axis) as a function of bulk concentration (x-axis). C) Total protein partitioning into assemblies (y-axis) as a function of bulk concentration (x-axis). D) Diameter of reflectin A1 assembly sizes as a function of bulk protein concentration. Data points for (B,C) represent 3 experimental replicates with 3 measurement replicates each. Error bars represent ± 1 S.D. For (D), 3 technical replicates of 3 experimental replicates each are shown as individual lines. Y-axis is volume percent and x-axis diameter in nm.

Reflectin A1 assembly sizes decrease as C*_bulk_* is increased from 1-4 μM **(Figure 1D)**. Notably, the size distribution of assemblies narrows as C*_bulk_* is increased from 1-10 μM **(Figure 1D)**. Assembly sizes range from 70-500 nm diam. for C*_bulk_* of 1 μM. At C*_bulk_* = 2 μM, size distributions are bimodal with a larger proportion by volume of assemblies in the smaller size distribution **(Figure 1D)**. As C*_bulk_* is further increased to 10 μM, this bimodality decreases **(Figure 1D)**. Reflectin A1 assembly size as a function of concentration mirrors the dependence of C*_dilute_* on C*_bulk_*. Partitioning of reflectin A1 into assemblies for C*_bulk_* ≤ 2 μM varies from C*_bulk_* ≥ 4 μM which is relatively constant **(Figure 1C)**. Assembly sizes for C*_bulk_* ≤ 2 μM vary from assembly sizes for C*_bulk_* ≥ 4 μM which is then relatively constant **(Figure 1D)**. Importantly, reflectin A1 assembly sizes do not increase with increasing protein concentration over the range analyzed in these conditions.

### Forster Resonant Energy Transfer (FRET) Reveals Reflectin A1 Assemblies Dynamically Exchange Protein Molecules With the Dilute Phase

The following experiment was performed to determine whether transport of protein occurs bidirectionally between reflectin assemblies or only unidirectionally as non-dynamic assemblies from monomers. Reflectin A1 assemblies labeled with both fluorescein (FRET donor) and rhodamine (FRET acceptor) were diluted with unlabeled A1 assemblies **(Figure 2A)**. A decrease in FRET would represent a decrease in the concentration of fluorophores within the protein assemblies, which would necessitate influx of unlabeled and efflux of fluorescently labeled reflectin A1. After diluting fluorescently labeled reflectin A1 with unlabeled A1 immediately after assembly formation, a decrease in FRET was observed in emission spectra **(Figure 2B)**. Replotting FRET as a function of time demonstrates that the dynamic exchange of reflectin between assemblies slows as the assemblies age **(Figure 2C)**. Reflectin A1 assemblies diluted immediately after formation are completely dynamic as FRET decreases to the positive control value **(Figure 2C)**. Relative to the range of the positive control (fully dynamic) to the negative control (fully arrested), 75.6% of the population of assemblies exhibits dynamic exchange after 1 min **(Figure 2C)**, while at 2 h 48% of the population remains dynamic, with only 19% still dynamically exchanging after 37 h. Reflectin A1 assembly sizes increased over the same time scale as FRET measurements of dynamic exchange **(Figure 2D)**. Initial assembly diameter was 24 ± 1.6 nm, and the assemblies grew to 33.9 ± 0.35 nm diam. after 24 h. The change in assembly size mirrors the decrease in dynamic exchange monitored by FRET **(Figure 2C,D)**. This is consistent with an arrest of Ostwald ripening, a mechanism by which larger condensates increase in size by exchange of particles through from the dilute phase (44).

**Figure 2.**
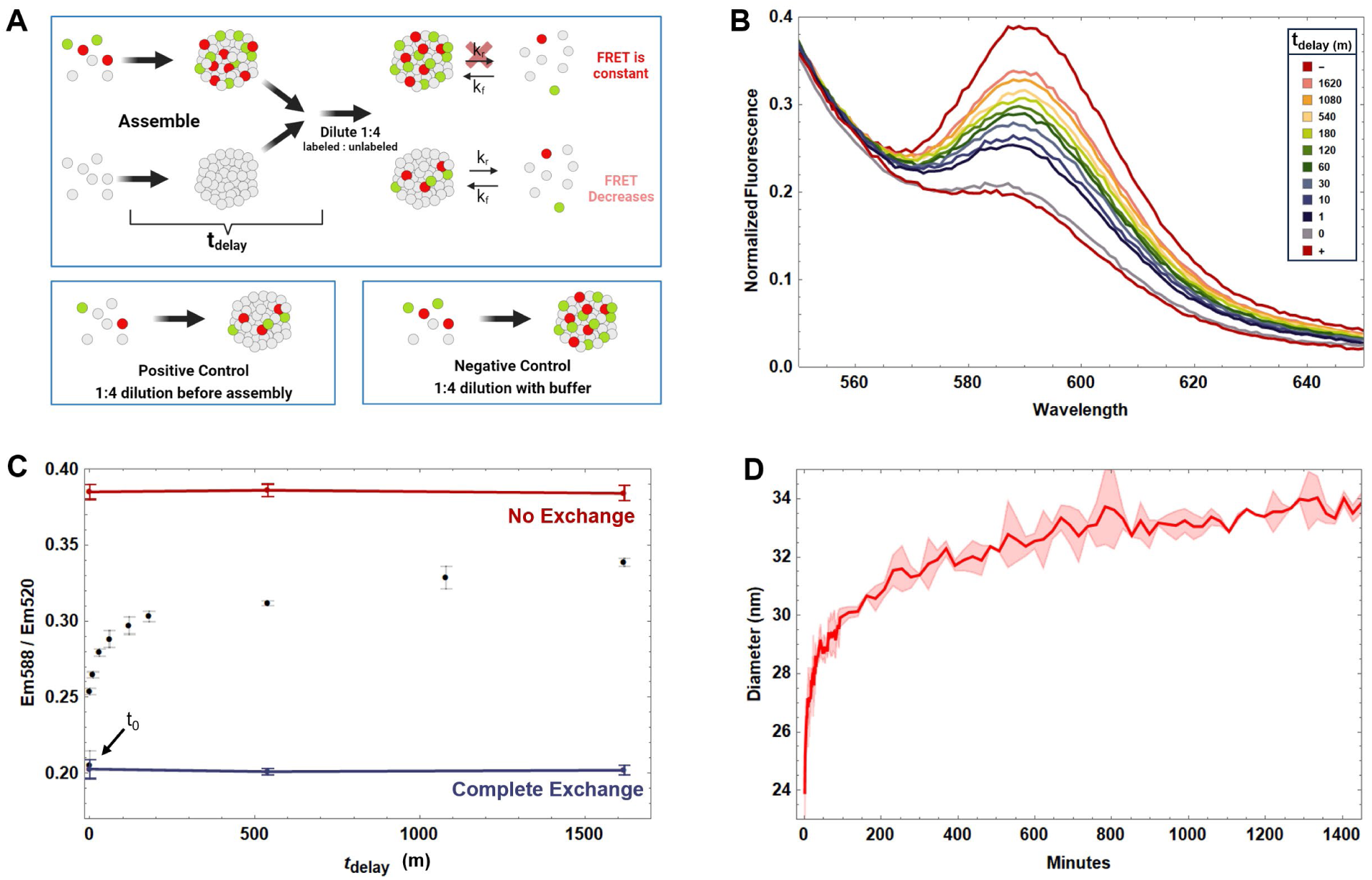
FRET assay monitors dynamic exchange of reflectin A1 assemblies as a function of time. A) Design of FRET dilution experiment. Mixtures of 100 µM reflectin A1 containing 5% fluorescein-labeled single cysteine mutant C232S (A1 C199-F) and 5% rhodamine sulfate-labeled C232S (A1 C199-R) were driven to assemble by dilution into MOPS buffer (pH 7, 25 mM). Simultaneously, unlabeled reflectin was driven to assembly under the same conditions. As a function of time post-assembly (t_delay_), labeled assemblies were diluted 1:4 with unlabeled assemblies then incubated for 24 h. Samples were excited with 488 nm wavelength laser corresponding to the absorption maximum of the donor fluorophore fluorescein and emission spectra from 510-700 nm recorded. For the negative control, fluorescently labeled assemblies were diluted 1:4 with buffer only. Reflectin A1 solutions containing 5% A1 C199-R and 5% A1 C199-F were diluted 1:4 with unlabeled reflectin prior to assembly for the positive control. B) Emission spectra normalized to 520 nm (Em_max_ of fluorescein) as a function of t_delay_, the time between assembly and mixing of labeled and unlabeled assemblies. C) FRET emission at 588 nm normalized to 520 nm and plotted as a function of t_delay_. The negative control represents zero dynamic exchange (red line) and the positive control represents complete dynamic exchange (blue line). D) In parallel, assemblies of reflectin A1 were formed by dilution of same protein stock into MOPS buffer (pH 7, 25 mM) and sizes determined by dynamic light scattering. For DLS, data points represent 3 experimental replicates. Error bars (C) and error bands (D) represent ± 1 S.D.

### Increasing Salt Concentration Drives Liquid-Liquid Phase Separation of Reflectin A1

The dilution of 100 µM reflectin A1 containing 5% single cysteine mutant A1 C232S covalently labeled with fluorescein (referred to as A1 C199-F to denote the amino acid location of fluorescein) into acetic acid buffer (pH 4, 25 mM) in 250 mM NaCl forms liquid condensates that wet untreated glass coverslips **(Figure 3A)**. Wetting is prevented using polyethylene glycol (PEG) passivated coverslips **(Figure 3B),** causing the condensates to remain spherical and undergo fusion with relaxation to sphericity upon contact **(Figure 3C)**. The rate of change of aspect ratio of newly fused droplets fits well with an equation of exponential decay that is commonly used to characterize liquid droplet dynamics (45, 46) **(Figure 3C)** yielding a relaxation time *τ* of 1.95 ± 1.16 sec. Surface wetting and relaxation of newly fused droplets indicate that the condensates exhibit surface tension which is characteristic of a liquid (19, 44).

**Figure 3.**
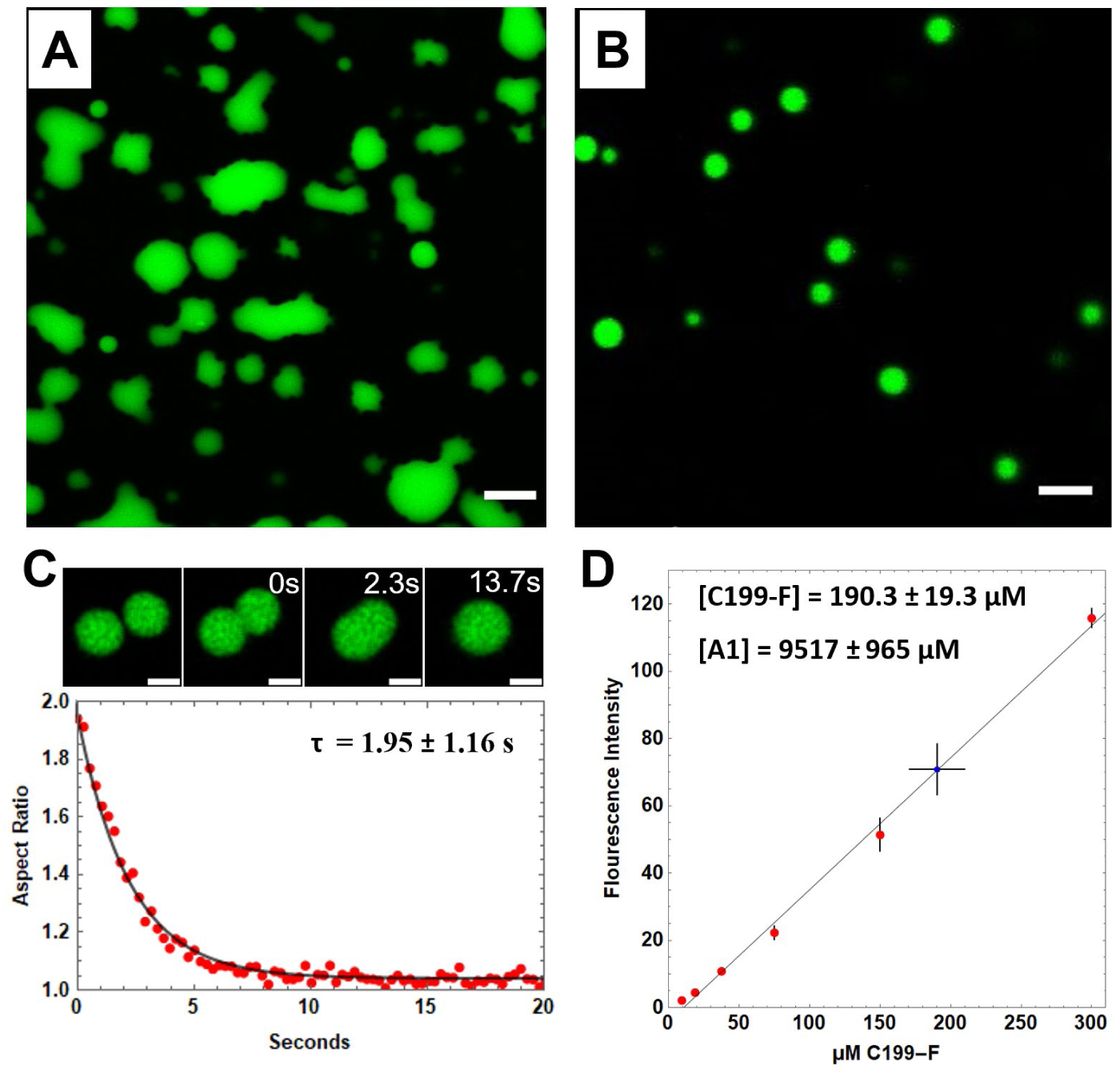
Confocal microscopy of fluorescently labeled reflectin A1 droplets. 100 µM reflectin A1 containing 5% fluorescently labeled A1 C199-F was diluted to a final protein concentration of 4 µM in acetic acid buffer (pH 4, 25 mM) in 250 mM NaCl. A) Reflectin liquid-like condensates wet untreated glass coverslips. B) Condensates form spherical droplets on PEG-passivated coverslips. C) Condensates fuse and relax to sphericity. The change in aspect as a function of time fits to an equation of exponential decay yielding the relaxation time τ. D) Reflectin A1 inter-droplet, or dense phase, protein concentration is 9517 ± 965 µM, or 416 ± 42 mg/mL. Monomeric standards (red) of 100% A1 C199-F in acetic acid buffer (pH 4, 25mM) were used to relate fluorescence intensity to concentration. Intensities of droplets of 2% C199-F were averaged (blue), (n=147 droplets from 5 experiments). Error bars = ± 1 S.D. Scale bars = 5 µm.

The results described above show that under these conditions, reflectin A1 undergoes liquid-liquid phase separation to form protein-dense and protein-dilute liquid phases. Comparison of the mean fluorescence intensity of reflectin A1 containing 2% A1 C199-F in acetic acid buffer (pH 4 25 mM) in 250 mM NaCl to that of 100% A1 C199-F monomeric standards in acetic acid buffer (pH 4 25 mM) determined the dense phase protein concentration to be 9,517 ± 965 µM, or 416 ± 42 mg/mL **(Figure 3D)**. This concentration is within the range of dense phase protein concentrations reported for many other liquid protein condensates (47–49) and is similar to reflectin protein concentration within the Bragg lamellae of *Doryteuthis opalescens* iridocytes, which was previously estimated to be 381 mg/mL (12). Thus, compared to the assemblies previously observed in low salt conditions, reflectin condensates may represent a more physiologically relevant biophysical model for investigating reflectin protein interactions.

### Reflectin A1 Phase Diagram Shows Reciprocal Relationship Between NaCl Concentration and Protein Net Charge Density

To investigate the role of protein net charge density and salt concentration on reflectin A1 phase transitions, as well as the potential for the use of reflectin A1 in tunable biomaterials, we determined the phase diagram for reflectin A1 as a function of ionic strength and net charge density, which is defined by three species: monomer, assembly, and liquid condensate **(Figure 4A)**. DLS was used to determine the boundary between monomer and assembly using the previously established R_h_ of reflectin A1 monomer (17, 21, 23) **(Figure 4D)**. The boundary between reflectin A1 assemblies and liquid condensates was determined by confocal microscopy of reflectin A1 containing 5% A1 C199-F. Droplets with the previously mentioned liquid characteristics (**Figure 4B)** were clearly distinguished from observable and sub-resolution assemblies **(Figure 4C)**. As the liquid phase boundary was approached by increasing ionic strength at a given pH (and its corresponding protein NCD), assemblies increased from sub-resolution to microscopically resolvable sizes **(Figure S5)**. Upon crossing the liquid phase boundary, protein partitioning into the dense phase increased markedly **(Figure S5)**. Additionally, the presence of slow modes in the DLS autocorrelation functions confirmed the observed microscopic distinctions between size-stable assemblies and liquid condensates at high NCD. Droplets with the previously mentioned liquid characteristics were detected microscopically upon dilution of reflectin A1 into 100 mM NaCl in acetic acid buffer (pH 4, 25 mM) **(Figure S6A)**, with DLS autocorrelation slow modes present and increasing with time **(Figure S6B),** which is characteristic of LLPS (41). In marked contrast, the same analyses in 90 mM NaCl in acetic acid buffer (pH 4 25 mM) failed to produce resolvable droplets **(Figure S6A)**, and the DLS autocorrelation functions did not change with time nor display slow modes **(Figure S6B)**.

**Figure 4.**
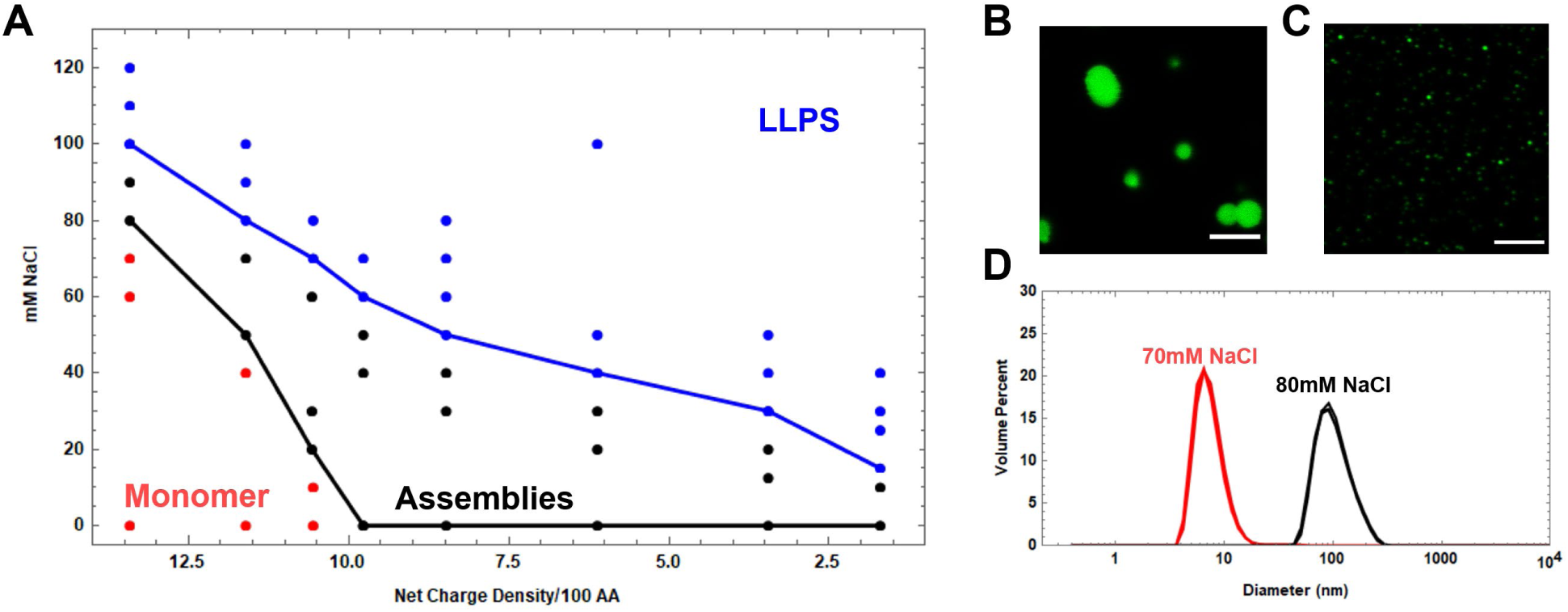
Phase diagram of reflectin A1 as a function of NaCl concentration and calculated protein net charge density. A) Reflectin A1 can exist as monomer (red) or assemblies (black), or liquid-like condensates (blue). B) Liquid-like droplets were distinguished from (C) light-resolvable assemblies by size discrepancy as well as surface wetting and droplet fusion. D) Assemblies were distinguished from monomers by using DLS to measure particle size distributions. 3 DLS replicates at each condition are shown cumulatively. At pH 4 in 70 mM NaCl, reflectin A1 exists as 7.9 ± 3.5 nm diam. monomers, while at pH 4 in 80 mM NaCl the protein forms 106 ± 39 nm diam. assemblies. Scale bars = 5 µm. At high pH, deprotonation of histidines decreases the protein net charge density per 100 amino acids. Net charge density is calculated from the pKa of each amino acid residue of reflectin A1 at each experimentally manipulated pH (11).

The decreasing slopes of both phase boundaries at high NCDs reveal that as the cationic charge of the protein is progressively neutralized (NCD is decreased), a progressively lower ionic strength is required to drive assembly or LLPS, consistent with the suggestion that pH titration and charge screening from salt act similarly (21). For lower protein NCDs (pH 5.5-8), reflectin A1 is not monomeric at any salt concentration in 25 mM buffers. For all protein NCDs tested, the assembly/LLPS boundary shares the same trend of decreasing protein NCD requiring decreasing ionic strength to drive LLPS **(Figure 4A)**. This trend at moderate salt concentrations, where Debye Lengths are similar to the distances of ionic interactions (43, 50), further supports the suggestion that ionic charge screening from salt drives both assembly and LLPS (51–53). In addition to increasing the ionic strength’s contribution to screening of repulsive charges (23), surface tension will also increase, thereby reducing protein solubility by increasing the strength of any hydrophobic drivers of assembly and LLPS.

### Salt Drives LLPS by Increasing Both the Hydrophobic Effect and Anionic Screening

To determine the possible role of the hydrophobic effect in driving reflectin A1 assembly and LLPS, the protein’s behaviors were compared in the presence and absence of 5% 1,6-hexanediol (1,6-HD). This water-soluble aliphatic alcohol lowers the surface tension of water (35) and has been shown to dissolve liquid protein condensates that are formed by the hydrophobic effect (35, 54–56). Turbidity measurements of reflectin A1 diluted into acetic acid buffer (pH 4, 25 mM) from 80 to 100 mM NaCl demonstrate that 5% 1,6-HD inhibited the salt-driven formation of large, light-scattering particles of reflectin A1 **(Figure 5A)**. In the absence of 1,6-HD, turbidity increased at 90 mM NaCl and was approximately 2.5 at 130 mM NaCl. In contrast, in the presence of 5% 1,6-HD, turbidity increased at 110 mM NaCl but reached only 1.5 **(Figure 5A)**. Similarly, while the dilution of reflectin A1 into 80 mM NaCl at pH 4 formed 52.7 ± 18.3 nm diameter assemblies, the inclusion of 5% 1,6-HD prevented this assembly **(Figure 5B)**. In 100 mM NaCl at pH 4, reflectin A1 formed liquid droplets, but the inclusion of 5% 1,6-HD abrogated LLPS and droplets were not observed **(Figure 5C)**. Further, DLS analyses reveal slow modes in the autocorrelation function at 100 mM NaCl at pH 4 which are no longer seen after the inclusion of 5% 1,6-HD **(Figure S4C)**.

**Figure 5.**
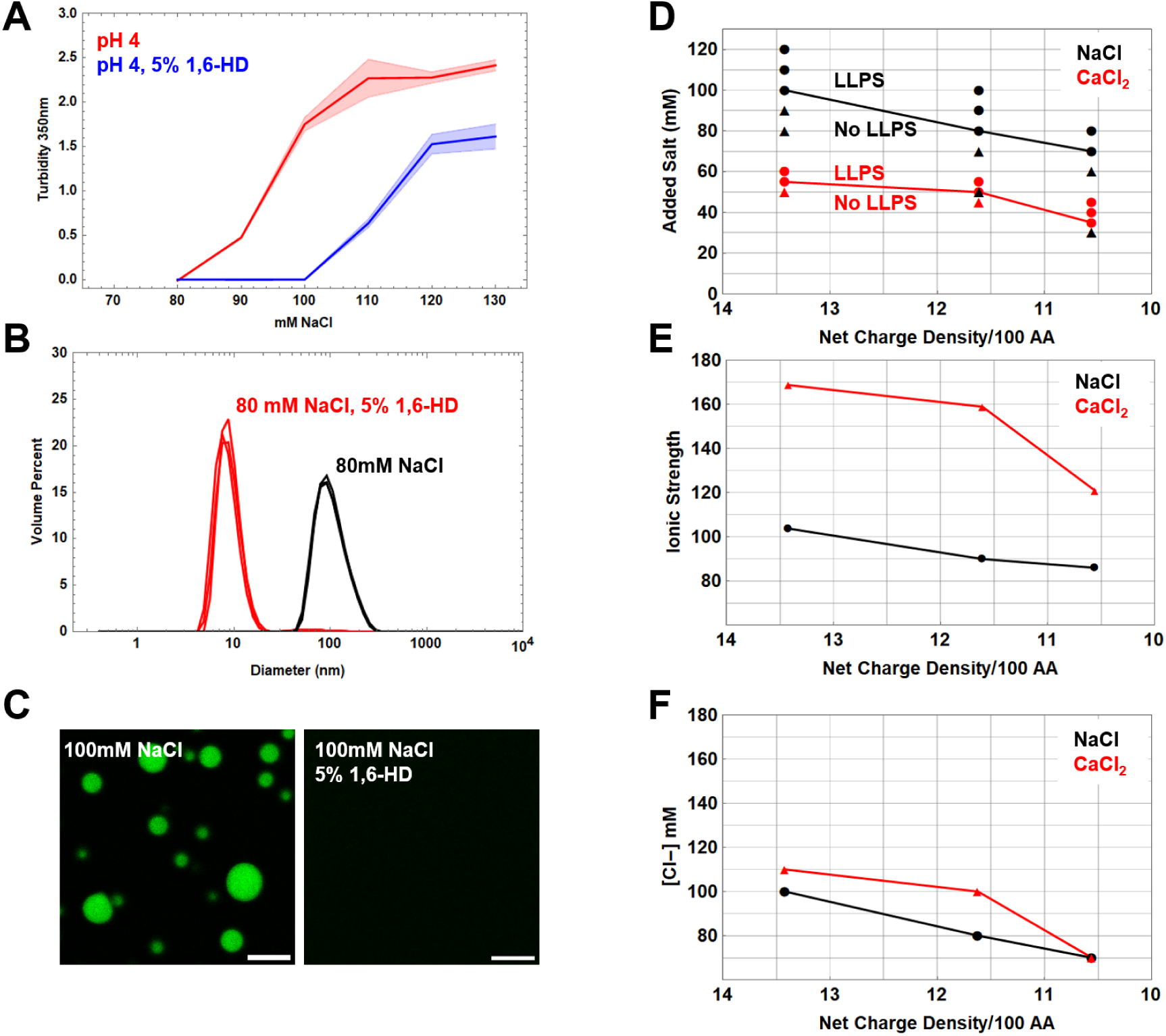
Effects of 1,6-hexanediol and anion concentration on reflectin A1 phase transitions. A) Reflectin A1 was diluted into 80-130 mM NaCl (red) with 5% 1,6-HD (blue) and turbidity measured at 350 nm. B) DLS of dilution of reflectin A1 into pH 4 80 mM NaCl (black) results in 106 ± 39 nm diam. assemblies and dilution into pH 4 80 mM NaCl, 5% 1,6-HD with yields monomers of 9.0 ± 2.6nm diam. C) Dilution of reflectin A1 into pH 4 100 mM NaCl forms droplets but the inclusion of 5% 1,6-hexanediol prevents droplet formation. Turbidity measurements are shown as the average of 3 experimental replicates with 3 measurement replicates each; error bands represent ± 1 S.D. The same numbers of DLS replicates are shown cumulatively, and images in (C) are characteristic of 3 experimental replicates. Scale bar = 10 µm. D) Liquid phase boundaries determined presence or absence of droplets upon addition of NaCl (Black) or CaCl_2_ (Red). E) Same data as (D) displayed as a function of ionic strength and (F) Cl^−^ concentration. X-axis for (D,E,F) is calculated protein net charge density using the method described in Figure 4. Data for NaCl addition is the same as that used for Figure 4. For all experiments 2 μL of 100 μM reflectin A1 in acetic acid buffer (pH 4 25 mM) was diluted into 48 μL of respective buffer.

These results indicate that increasing ionic strength drives reflectin A1 assembly and LLPS in part by increasing the strength of the hydrophobic effect. This would be expected because of the large gain in entropy typically resulting from burying hydrophobic side chains away from water (57–59) and LLPS is primarily a segregative process (19) driven by differences in polymer and solvent polarity.

It has been previously demonstrated that the sizes of reflectin A1 assemblies formed by addition of salt are well predicted by the concentration of anions (23). To investigate if this relationship is also true for salt-induced LLPS of reflectin A1, the liquid phase boundary of reflectin A1 was determined as a function of CaCl_2_ concentration in addition to NaCl concentration for pH values 4-5 **(Figure 5D)**. The liquid phase boundaries as a function of anionic concentration **(Figure 5E)** differ by 9.6% for pH 4, 21% for pH 4.5, and are identical for pH 5. In comparison, the liquid phase boundaries as a function of ionic strength **(Figure 5F)** vary by 62% for pH 4, 48.9% for pH 4.5, and 20.4% for pH 5. Therefore, the concentration of anionic species better describes the liquid phase boundary determined by addition of NaCl and CaCl_2_ than does ionic strength, which strongly suggests that electrostatic screening of cationic Coulombic repulsion within and between reflectin A1 molecules contributes to the drive of reflectin A1 to undergo LLPS. Screening of long-range repulsion at high protein NCDs would allow the short-range attractive forces to drive reflectin A1 folding, assembly and LLPS. We conclude that increasing salt concentration and reducing protein NCD drive reflectin A1 phase transitions by the same physical processes: both reduce protein solubility as well as neutralize protein net charge density.

### Reflectin A1 Condensates Demonstrate Liquid Properties up to 96 Hours

FRAP was used to investigate the aging of reflectin A1 droplet dynamics **(Figure 6A)** over time scales that far exceed *in vivo* reflectin activation (12), but are relevant for the use of reflectin A1 as a tunable biomaterial. The percent fluorescence recovery from FRAP of reflectin A1 condensates, which represents the proportion of the liquid-like population of the dense phase, in pH 4 25 mM acetic acid 250mM NaCl decreased significantly from 0-24 h, does not change significantly from 24-72 hours, and then decreased over from 72-96 h. **(Figure 6B)**. Remarkably, *τ* did not change significantly from fresh droplets to 72 h, but then increased drastically from 72-96 h **(Figure 6C)**. Reflectin A1 droplets showed no visible changes in morphology such as jaggedness or fibrilization for the duration of the experiment **(Figure 6D)**. Interestingly, despite the stability of τ in these conditions, droplet size did not change significantly over 96 h suggesting that Ostwald ripening was arrested **(Figure 6D)** (60–62). The time-dependent decrease in both the liquid-like proportion of proteins and the diffusivity of this population within reflectin A1 droplets may be due to an increase in non-covalent interactions and/or progressive dehydration (63, 64).

**Figure 6.**
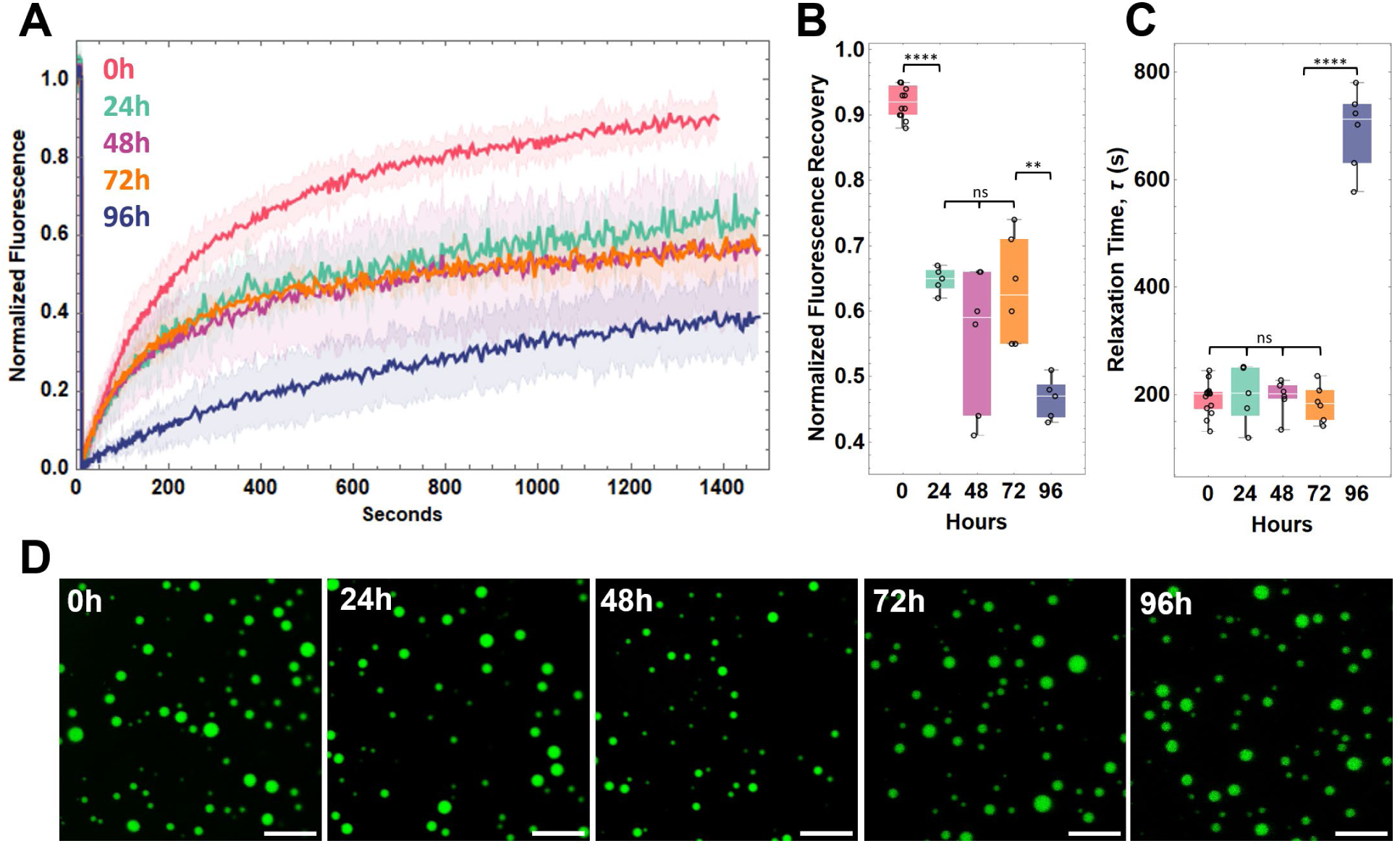
Time-dependent diffusivity of reflectin A1 condensates. A) FRAP of reflectin A1 liquid-like condensates in pH 4 25 mM acetic acid 250 mM NaCl for 0-4 (red), 24 (green), 48 (purple), 72 (orange), and 96 hours (blue). B) Data were corrected for photobleaching and fully normalized fluorescence intensity of bleach spots plotted as a function of time. C) Fitting of individual experiments to an exponential decay equation yields the characteristic relaxation time τ. Statistical significance determined by one-way ANOVA testing. Box and whisker plots display the median (center line), 2nd and 3rd quartile (solid box) and the complete range (black fences) of the data. D) Droplet morphologies for 0-96 hours. N=12 for 0h, N=5 for 24h, N= 6 for 48h, N=6 for 72h, N=6 for 96h.

### Net Charge Density and Ionic Strength Tune Reflectin Liquid Condensate Dynamics

Motivated by the potential use of reflectin A1 for tunable biomaterials, we used FRAP and droplet fusion dynamics to determine that the liquidity of reflectin A1 condensates is tuned by protein NCD. Videos of reflectin A1 condensates in 250mM NaCl at pH 4, 6, and 7 **(Figure 7A,B,C)** were used to determine the change in aspect ratio of newly fused droplets as a function of time. Fitting these data to an equation of exponential decay **(Figure 7D,E,F)** reveals that relaxation time *τ* increased from 1.95 ± 1.16s at pH 4 to 36.95 ± 29.32s at pH 6, and ultimately to 84.86 ± 22.77s at pH 7 **(Figure 7G)**. Droplet fusion and relaxation rates increased by almost two orders of magnitude from pH 4 to pH 7, demonstrating that liquidity (46) of reflectin A1 condensates decreases as protein NCD decreases.

**Figure 7.**
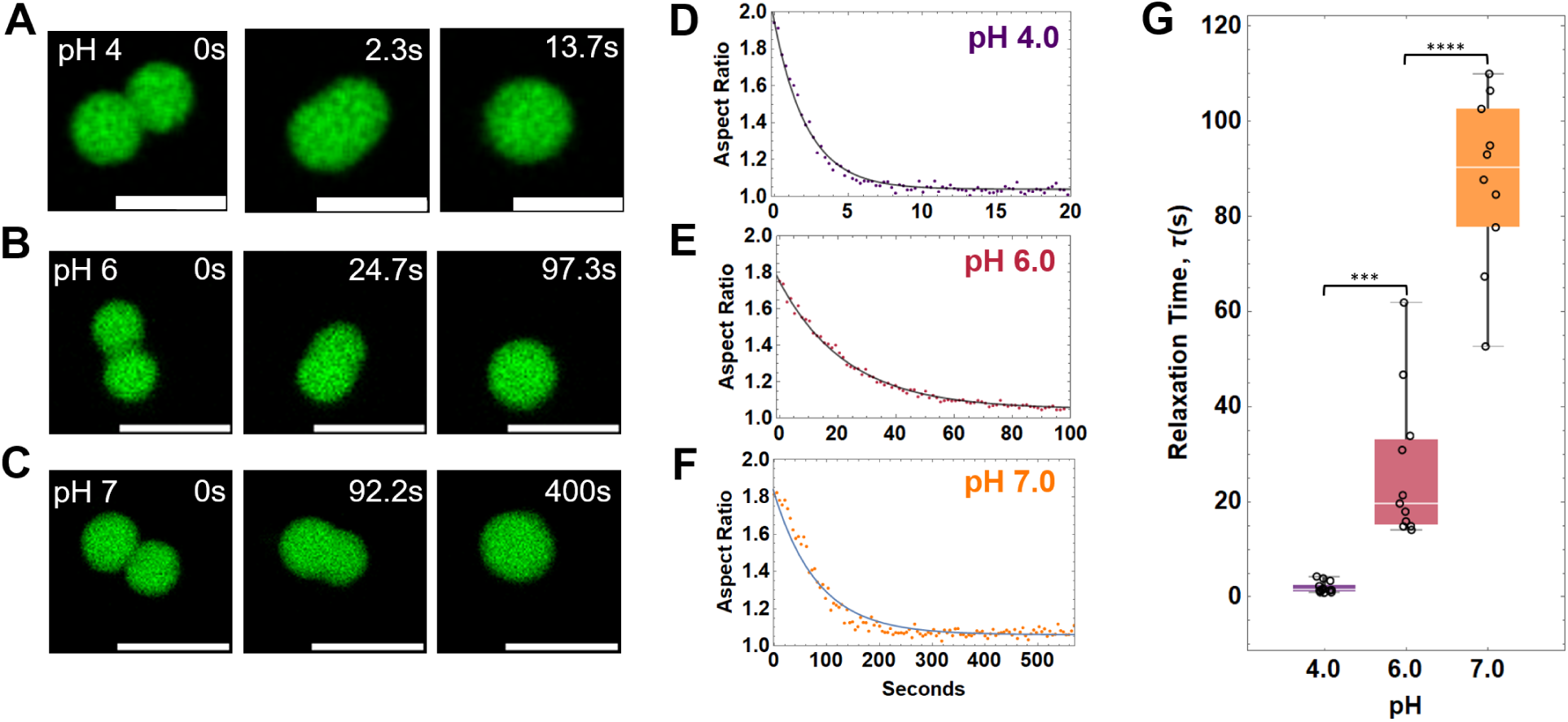
Quantification of reflectin A1 condensate fusion dynamics. Aspect ratios of newly fused droplets in 250 mM NaCl at pH 4, 6 and 7 were calculated from videos. Representative time series of droplet fusion and relaxation at (A) pH 4, (B) pH 6 and (C) pH 7. Aspect ratio as a function of time was fit to an exponential equation of decay to determine τ for (D) pH 4, (E) pH 6, and (F) pH 7. G) Changes in aspect ratio over time between each pH were found to be statistically significantly different using a one-way ANOVA test. Box and whisker plots display the median (center line), 2nd and 3rd quartile (solid box) and the complete range (black fences) of the data. N=23 for pH 4, n=12 for pH 6, and n=10 for pH 7. Scale bars = 5 µm.

FRAP analyses of reflectin A1 droplets in 250 mM NaCl at pH 4, 6 and 7 **(Figure 8A,B)** demonstrate a dependence of percent fluorescence recovery and *τ* on protein NCD. The proportion of mobile component of the droplets decreases as pH increases (protein NCD decreased) **(Figure 8C)**, indicating that the population of liquid-like protein molecules in the droplet is decreasing. Fluorescence recovery rates *τ* increase from 192.3 ± 31.1 s at pH 4 to 641.6 ± 220.2 s at pH 6, and at pH 7 the dynamics within reflectin condensates were too slow to determine accurate *τ* values **(Figure 8D)**. This dependence on protein NCD could be due to more rapid aging of non-covalent protein interactions at lower protein NCDs (higher pHs) than at higher protein NCDs (lower pHs), as more extensive protein-protein interactions would be expected for decreasing Coulombic repulsion at low protein NCDs. It also is possible that reflectin A1 condensates formed at low protein NCDs are less hydrated and more protein-dense than condensates formed at high protein NCDs, further contributing to the slowing of condensate dynamics. Reflectin A1 droplets showed slower and incomplete fluorescence recovery compared to many other *in vitro* and *in vivo* protein systems (33, 38, 45, 65–70).

**Figure 8.**
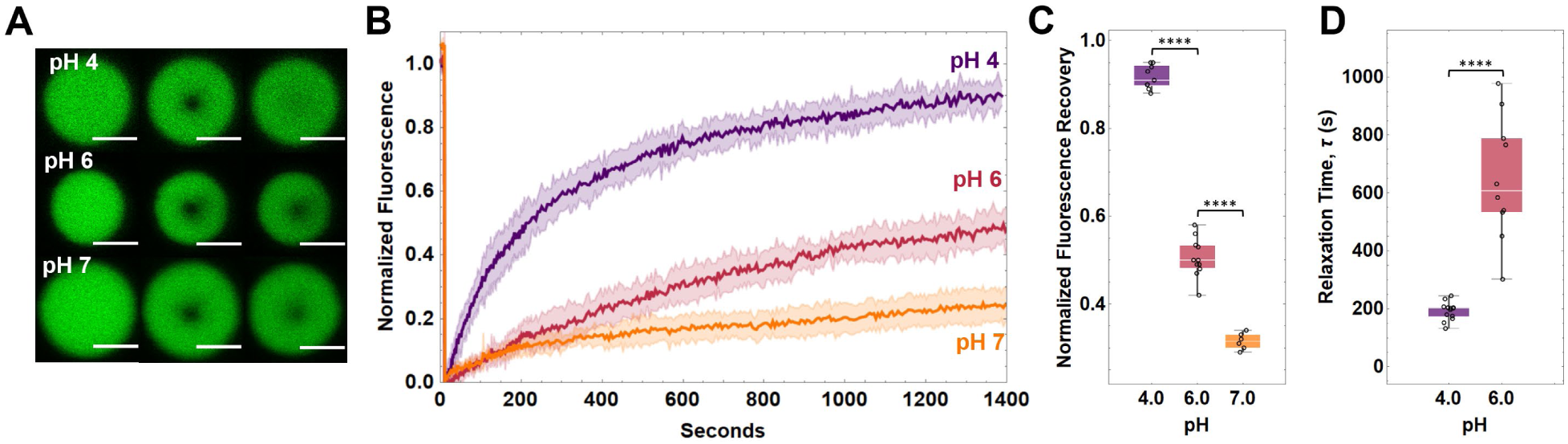
FRAP of reflectin A1 condensates at pH 4, 6 and 7 in 250 mM NaCl. A) characteristic images of droplets before, immediately and 1400s post photobleaching for pH 4,6, and 7. B) Data were corrected for photobleaching and fully normalized fluorescence intensity of bleach spots plotted as a function of time. C) Percent fluorescence recovery for pH 4, 6, and 7. D) Fitting of individual experiments to an exponential decay equation yields the characteristic relaxation time τ. τ for pH 4 and pH 6, as well as percent recovery for all pHs were found to be statistically significant using a one-way ANOVA test. Box and whisker plots display the median (center line), 2nd and 3rd quartile (solid box) and the complete range (black fences) of the data. Accurate τ values for pH 7 could not be obtained by fitting. N=12 for pH 4, n=11 for pH 6 and n=7 for pH 7. Error bands in FRAP plots represent ± 1 SD. Scale bars = 5 µm.

## Discussion

Reflectin A1’s net charge density (NCD) has been shown to finely control the protein’s assembly sizes, and it has been hypothesized that progressive neutralization of the protein’s excess cationic charge, reducing Coulombic repulsion both within and between reflectin molecules, mediates this relationship (11), both physiologically in response to neuronally activated phosphorylation (6, 10) and *in vitro* by various surrogates (11, 21, 22). Here, we extend these results to demonstrate that further reduction of Coulombic repulsion by ionic screening drives reflectin A1 to undergo LLPS into liquid condensates. Notably, as salt concentration increases, assemblies increase in size **(Figure S5)** (23) and reflectin A1 assembly always precedes LLPS, suggesting that reflectin A1 assembly and LLPS are stages of the same physical path (44, 60). The dependence of the liquid phase boundary of reflectin A1 on the concentration of anions at high protein NCDs **(Figure 5)** supports the hypothesis that ionic screening of Coulombically repulsed cationic reflectin A1 allows for attractive forces (e.g., cation-pi, sulfur-pi, hydrophobic effect) to drive LLPS of reflectin A1. Importantly, this matches and extends the recent demonstration that salt induces reflectin A1 assembly by ionic screening (23), and both mechanisms are consistent with the findings that the extent of reduction of reflectin A1’s NCD by phosphorylation, pH titration, genetic engineering or electroreduction proportionally control reflectin A1 assembly sizes (11). The phase behavior of reflectin A1 can be compared with that of other proteins exhibiting both assembly and LLPS behavior (41, 71, 72), notably FUS cluster formation in a pre-percolation state of percolation-coupled phase separation. That behavior is not consistent with our observations of reflectin A1 assemblies, in which (i) 95-99% of bulk reflectin A1 protein is found in assemblies; (ii) reflectin A1 assemblies are relatively monodisperse and display a normal distribution of sizes as detected by DLS in our observations and previously (11, 17, 23); and (iii) reflectin A1 assembly sizes did not increase with increasing concentration for the range of 1-10 µM. Contrary to the behavior of reflectin A1, solutions of proteins FUS, TDP-43, BrD4, Sox2 and A11 undergo LLPS under low salt concentrations (<50 mM KCl), exist as a single phase at intermediate salt concentrations (125 mM-1.5 M KCl), and are prevented from undergoing LLPS by the disruption of attractive ionic protein interactions by electrostatic screening at intermediate salt concentrations (54). Hence, it is not without precedent that intermediate salt concentrations [concentrations where the Debye distance is similar to that of electrostatic interactions between amino acid side chains (43, 50)] can dictate liquid phase transitions of proteins by screening ionic interactions, albeit with reflectin it is the screening of repulsive electrostatic interactions that drives assembly and LLPS.

Both reflectin A1 assembly and LLPS are driven by increasing the relative strength of the hydrophobic effect. Previously, shifts in fluorescence spectra of the native tryptophans in reflectin A1 as well as hydrophobic binding of the fluorophore, ANS, demonstrated a net transfer of these fluorophores into a more hydrophobic environment upon reflectin A1 assembly (17). Although the primary sequence of reflectin A1 does not have any significant stretches of hydrophobic amino acids, computational analyses of hydrophobic moments, which are the net amphiphilicity of a given protein segment as a function of the angle between successive side chains (73), show the highly conserved RMs have maximal hydrophobic moments at side chain angles consistent with *α*-helical (100*°*) and *β*-sheet (160*°*) secondary structures (16), and these secondary structures emerge upon folding and assembly of reflectin A1 (11, 22). These results suggest that the emergent hydrophobicity upon secondary structure formation in the RMs may provide a major hydrophobic contribution to folding, assembly and LLPS of reflectin A1.

Thermodynamic equilibrium, as opposed to kinetic arrest or glassification, could define reflectin A1 assembly size by a trade-off between short range attractive forces and long-range repulsive forces between monomers (SALR) (74). Equilibrium protein clusters described by SALR models have been reported (75, 76) in which long range electrostatic repulsion competes with Van der Waals forces and the hydrophobic effect to control cluster size. The electrostatic repulsion between cationic reflectin A1 proteins can be described as a long-range repulsive force that, in continual balance with short range attractive forces (including those of hydrophobic, cation-pi and sulfur-pi interactions) define both the protein’s assembly sizes and phase transitions. Screening of Coulombic repulsion by added anions, shown to be proportional to equilibrium cluster sizes in both lysozyme and hemoglobin solutions (75–78), have been shown to control reflectin A1 assembly sizes similarly (23). In a model colloidal system, the formation of equilibrium clusters of lysozyme can be followed by a glass transition that arrests cluster dynamics (75), although this transition occurs much more rapidly than the observed decrease in reflectin A1 dynamics as monitored by FRET **(Figures 1, 2)**.

*In silico* modeling demonstrated that mesoscopic metastable clusters can be intermediates to the condensation of a bulk liquid phase, and that increased ionic screening lowers the repulsive energy barrier and allows bulk transition to two liquid phases (79). Thus, LLPS of reflectin A1 may represent such a state in which the electrostatic repulsive energy barrier is no longer sufficient to restrict assembly size. Excitingly, computational modeling of reflectin A1 assemblies as a function of protein NCD (via pH titration) well describes A1 assembly size distributions as SALR-dictated equilibrium clusters (40). A colloidal model was developed in which reflectin monomers interact via a short-range attractive and long-range repulsive pair potential (SALR), with parameters of the model calibrated with data from the experimental short-angle x-ray scattering (SAXS) analyses of reflectin A1 assemblies in which size was tuned by pH titration. This model predicted that reflectin A1 assemblies are dynamic equilibrium clusters, successfully predicting assembly sizes as a function of protein NCDthat were consistent with previous DLS observations (11, 40). Decreasing the modeled long-range repulsive force increased equilibrium cluster size until LLPS occurred. Further, this analysis predicts a phase diagram for reflectin A1 as a function of protein NCD that agrees with that presented here for the protein concentrations used in our experiments. However, because reflectin A1 phase behavior is likely influenced by the valency of interprotein interactions, a possible shortcoming of relating this behavior to that of simple colloidal systems is that these do not depend on such valency – unlike many other proteins that undergo physiologically relevant LLPS (19, 44, 60, 74–80).

The complex phase behavior of reflectin A1 adds to our understanding of the possible states of matter in the Bragg lamellae in the tunable iridocytes of *D. opalescens. In vivo*, iridocyte activation and change in reflectance in response to acetylcholine occurs on the timescale of 10s of seconds (5, 10) and our FRET observations show little decrease in the rate or extent of dynamic exchange between assemblies over the same time **(Figure 2)**. This demonstrates that reflectin A1 assemblies can remain dynamic over the physiologically relevant time. The protein concentration in reflectin condensates was found to be 416 ± 42 mg/mL **(Figure 3D)**, similar to the previous estimate of the *in vivo* lamellar protein concentration of 381 mg/mL (12). Our finding that progressive neutralization of reflectin protein net charge density drives a phase transition from multimeric assemblies to a dense liquid phase parallels observations of the reflectin-containing iridocytes and leucophores in the living tissue, and observations of reflectin’s behavior in the isolated Bragg lamellae. Upon acetylcholine activation, reflectin in the iridocyte Bragg lamellae and leucophore vesicles undergoes a phase transition from 10-50 nm diameter particles into a dense liquid (5, 13, 39). Bragg lamellae isolated from activated iridocytes of the Loliginid squid *Lollinguncula brevis* showed liquid properties such as surface wetting and deformability, in contrast to those isolated from nonactivated iridocytes (39).

Further, expression of reflectin A1 from *D. pealeii* (81) and individual expression of reflectins A1, A2, B and C from *D. opalescens*, both in HEK 293T cells, yielded intracellular spherical puncta consistent with liquid-liquid phase separation (82). In tunable iridocytes, a liquid phase transition would maximize the lamellar protein density relative to a solution of assemblies and therefore enhance the refractive index contrast resulting from activation, while maintaining a dynamic environment essential for the controlling activities of reflectin-specific kinases and phosphatases. Decreasing the protein’s NCD by phosphorylation could provide a mechanism for tuning lamellar dehydration and the consequent photonic behavior (7, 8, 12) by controlling the dense phase protein concentration. Further, Bragg lamellae in tunable iridocytes from *D. opalescens* are enriched in reflectins B and C and relatively low in reflectin A2 compared to non-tunable iridocytes from the same species (6). It remains to be seen if reflectins A2, B and C also undergo LLPS in protein NCD-dependent manner, and if these species affect the liquid phase boundary and condensate properties of reflectin A1. Nonetheless, the finding that reflectin A1 can be driven to liquid-liquid phase separation by changing protein NCD and ionic strength, and that the liquid properties of these condensates is controlled similarly, illuminates a significant extension of both our understanding of Bragg lamellar condensation and the remarkable tunability of the reflectin system.

## Experimental Procedures

### Protein Expression and Purification

Reflectins A1, A1-C232S, and A1-6E were purified as previously described (11). Codon-optimized sequences of reflectins A1 and A1-C232S, were cloned into pj411 plasmids. Proteins were expressed in Rosetta 2 (DE3) cells grown in 1 or 2 liter LB cultures at 37°C from plated and sequenced transformants in the presence of 50 mg/mL kanamycin. Expression was induced at OD_600_ of 0.6-0.7 with 5 mM IPTG. After 16 h, cultures were pelleted by centrifugation and frozen at −80°C. Reflectins were expressed in inclusion bodies, which were purified using BugBuster medium (Novagen, Inc., Madison, WI) per manufacturer protocol, then resolubilized in 8 M urea 5% acetic acid. Reflectins were purified using cation exchange with a 10 mL Hitrap cation-exchange column (Cytiva, Marlborough, MA) and eluted using a step gradient (0% to 7% to 10%) of 5% acetic acid, 6 M guanidinium chloride. Reflectins eluted at 10% eluting buffer. Purity of collected fractions was determined by A260/A280 and SDS-PAGE, which were then pooled and concentrated. Concentrated reflectin was loaded onto reverse-phase HPLC Xbridge 4.6 mL C4 column (Waters, Milford, MA) equilibrated with 10% acetonitrile with 0.1% trifluoroacetic acid and eluted over a gradient of 100% acetonitrile 0.1% TFA. Fractions were frozen at −80°C or shell frozen using an ethanol and dry ice bath, lyophilized, and stored at −80°C until solubilization. Purity was determined by SDS-PAGE and A260/A280. Lyophilized reflectin was solubilized using 0.22 μm-filtered acetic acid buffer (pH 4 25 mM) and dialyzed using 2 changes (12 hours each) of 1000X sample volume of the same buffer at 4°C. Protein concentration was calculated using absorbance at 280 nm and molar extinction coefficients for each protein (reflectin A1 : 120685 M^−1^cm^−1^, reflectin A1 C232S : 120560 M^−1^cm^−1^). Protein stock was centrifuged at 18,000 X g for 15 minutes at 4°C prior to use in all assays. Protein stocks were stored at 4°C between uses.

### Bradford Assay

0.5, 1, 2, 3, 4, and 54 µL of 100 µM reflectin A1 in acetic acid buffer (pH 4 25 mM acetic acid) were driven to assembly by dilution into freshly 0.22 μm-filtered MOPS buffer (pH 7.0 25 mM) to a total volume of 50 μL and final protein concentrations of 1, 2, 4, 6, 8, and 10 µM, then incubated at 20° C for five days. Samples were then centrifuged at 20°C, 20,000 X g for 6 hours and a small portion of supernatant immediately removed, combined with Bradford dye and absorbances at 595 nm recorded after 30 minutes. A serial dilution of BSA standard was used to create a standard curve. For the Bradford assay, experimental data points represent averages of three experimental replicants with three measurements each. In parallel, centrifugation was omitted, and sizes were determined by dynamic light scattering (DLS) using a Malvern Zetasizer Nano. All DLS measurements were replicated with three individual experiments, each with three technical replicates spanning 5 minutes.

### Fluorescent Labeling

Reflectin A1-C232S was covalently labeled using cysteine specific fluorescein- or rhodamine-methanethiosulfonate (FMTS or MTSR). 50 µM protein was incubated with 500 µM FMTS or MTSR in acetic acid buffer (pH 4, 25 mM) with gentle orbital stirring for 4 hours RT then overnight at 4° C. Labeled reflectins were concentrated using 10K MWCO Amicon spin filters, and excess label removed using the previously described HPLC method (11). Labeling efficiencies for FMTS and RMTS were 100%.

### FRET

For FRET dilution experiments, 100 µM Reflectin A1 and 100 µM reflectin A1 containing 5% C199-F and 5% C199-R were separately but simultaneously diluted into MOPS buffer (pH 7, 25 mM) to final protein concentrations of 4 µM. Fluorescently labeled reflectin A1 assemblies were then diluted 1:4 with unlabeled reflectin A1 assemblies after the designated time t_delay_ and incubated in protein LoBind tubes at 20° C for 24 hours before measurements. Using a Cary Eclipse fluorescence spectrometer (Agilent Technologies, Australia), samples were excited at 488 nm and emission spectra recorded from 510-700 nm. Each data point is the average of three experimental replicates. For the positive control, solutions of 100 μM A1 and 100 μM A1 containing 5% each of C199-R and C199-F in acetic acid buffer (pH 4, 25 mM) were mixed, then assembled by dilution into MOPS buffer (pH 7, 25 mM). For the negative control fluorescently labeled assemblies were formed as per the experimentals but diluted 1:5 with MOPS buffer (pH 7, 25 mM). All spectra were normalized to 520 nm (Em_max_ of fluorescein).

For mixing of assemblies separately labeled with donor and acceptor fluorophores, 100 µM reflectin A1 containing 5% C199-F and 100 µM reflectin A1 containing 5% C199-R were separately but simultaneously driven to assembly by dilution into MOPS buffer (pH 7, 25 mM) to final protein concentrations of 4 µM. After a designated time, fluorescein-labeled and rhodamine-labeled A1 assemblies were mixed and treated identically to samples in the dilution experiment. For the positive control a solution of 100 µM reflectin containing both 5% C199-F and 5% C199-R was driven to assembly. The negative control mixed assemblies containing 5% C199-R with assemblies containing 5% C199-F 24 hours after assembly and measured immediately upon mixing. All buffers were freshly 0.22 μm-filtered before use.

### Phase Diagrams

To determine the liquid-liquid phase separation boundary, 0.6 µL 100 µM reflectin containing 5% fluorescein labeled protein (A1 C199-F) in acetic acid buffer (pH 4 25mM acetic acid) were diluted into 9.4 µL of respective buffers to a final protein concentration of 4 µM, tap mixed, and incubated for 10 minutes using 0.5 mL protein LoBind tubes. The solutions were pipetted onto cleaned glass coverslips and imaged immediately after deposition using Leica SP8 resonant confocal microscope with 63X objective (NRI-MCDB Microscopy Facility). To determine the boundary for transition from monomer to assembly, 2 µL of 100 µM reflectin in acetic acid buffer (pH 4 25mM acetic acid) were diluted into 48 µL respective buffer to a final protein concentration of 4 µM and incubated as above. Sizes were determined by dynamic light scattering (DLS) using a Malvern Zetasizer Nano. All DLS measurements were replicated with three individual experiments with three technical replicates spanning 15 minutes each. All buffers were freshly 0.22 μm-filtered before use.

### Turbidity Assays

2 µL of 100 µM reflectin in acetic acid buffer (pH 4 25mM acetic acid) were diluted into 48 µL of respective buffer in the measurement cuvette and tap mixed. After 5 minutes of incubation at 25° C, 3 replicant measurements were recorded. This was repeated for a total of 3 experimental replicates.

### Glass Coverslip Passivation

For FRAP and droplet fusion analyses, glass coverslips were passivated to prevent interactions between reflectin and the glass surface. Glass slides and coverslips were sonicated in near-boiling 1% Hellmanex cleaning solution for 15 minutes. After thorough rinsing with Milli-Q water, glass slides were dried with CO_2_ and stored with Drierite desiccant. Coverslips were immersed in 0.1 M NaOH for 15 minutes, thoroughly rinsed with Milli-Q water, then dried with CO_2_. They were then heated to 100° C in closed glass Petri dishes containing Drierite for 10 minutes, after which 20 µL of 1% methoxy-PEG silane was pipetted onto each 18X18 mm coverslip and incubated for 20 minutes at 100° C. After allowing to cool, coverslips were thoroughly rinsed, sonicated in Milli-Q water for 10 minutes, and rinsed again. After drying with CO_2_, glass chamber microscope slides were constructed using two strips of Scotch double sided tape to secure the coverslip against the slide.

### FRAP Experiments and Analysis

100 μM reflectin A1 containing 5% C199-Fluorescein was diluted into freshly 0.22 μm-filtered buffers, then quickly pipetted into PEG-passivated chamber slides which were sealed with fast-drying clear nail polish. Using the FRAP module in a Leica SP8 scanning confocal microscope, a 1 μm diameter circular ROI (“point ROI” in LASX software) was selectively photobleached at 45-50% laser power for 150-200ms. The bleached ROI was monitored for 10 s before bleaching, and for 25 m post bleaching. For data analysis, videos were stabilized in ImageJ using the StackReg plugin and the fluorescence intensity of the bleached region, an unbleached region in the droplet, and background were used to correct for photobleaching:

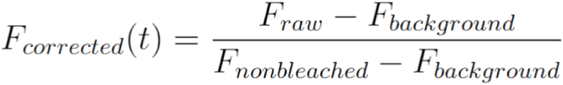

then fully normalized using the equation:

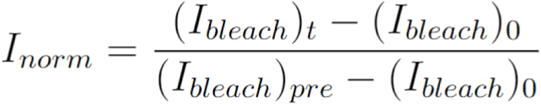

where (I*_bleach_*)*_t_* is the intensity of the photobleached region at time t (I*_bleach_*)_0_ is the intensity of the photobleached region at the time of photobleaching (I*_bleach_*)*_pre_* is the intensity of the photobleached region prior to photobleaching. To determine the characteristic relaxation time *τ* data was fit to the equation:

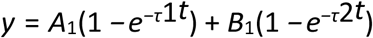

using the NonLinearRegression function in Mathematica.

### Droplet Fusion Analysis

Reflectin protein and slides were prepared as described for FRAP, and were imaged immediately after chamber slide was prepared. Only fusion events of equally sized droplets were analyzed. Using Fiji ImageJ software, the threshold function was used to create binary videos. Aspect ratios were calculated by using the “Fit Ellipse” measurement in “Analyze Particles”. Using the NonLinearRegression function in Wolfram Mathematica, the change in aspect ratio over time was fit to the equation:

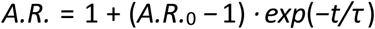

where A.R._0_ is the initial aspect ratio, t is the time, and *τ* is the characteristic relaxation time.

## Acknowledgements

Research was supported by the Institute for Collaborative Biotechnologies through grants W911NF-19-2-0026 and W911NF-23-1-0330 from the U.S. Army Research Office. The content of the information does not necessarily reflect the position or the policy of the Government, and no official endorsement should be inferred. We gratefully acknowledge the use of the facilities of UCSB’s NRI Microscopy Center and Ben Lopez for his guidance.

## List of Abbreviations

1,6-HD: 1,6-hexanediol
DLS: Dynamic Light Scattering
FRAP: Fluorescence Recovery After Photobleaching
FRET: Fluorescence Resonant Energy Transfer
IPTG: Isopropyl β- d-1-thiogalactopyranoside
LLPS: Liquid Liquid Phase Separation
MOPS: 3-(N-morpholino)propanesulfonic acid
NaCl: Sodium Chloride
NCD: Net Charge Density
RM: Reflectin Repeat Motif
RM_n_: N-Terminal Repeat Motif
SDS PAGE: Sodium Dodecyl Sulfate Polyacrylamide Gel Electrophoresis

## Supplemental Information

**Figure S1.**
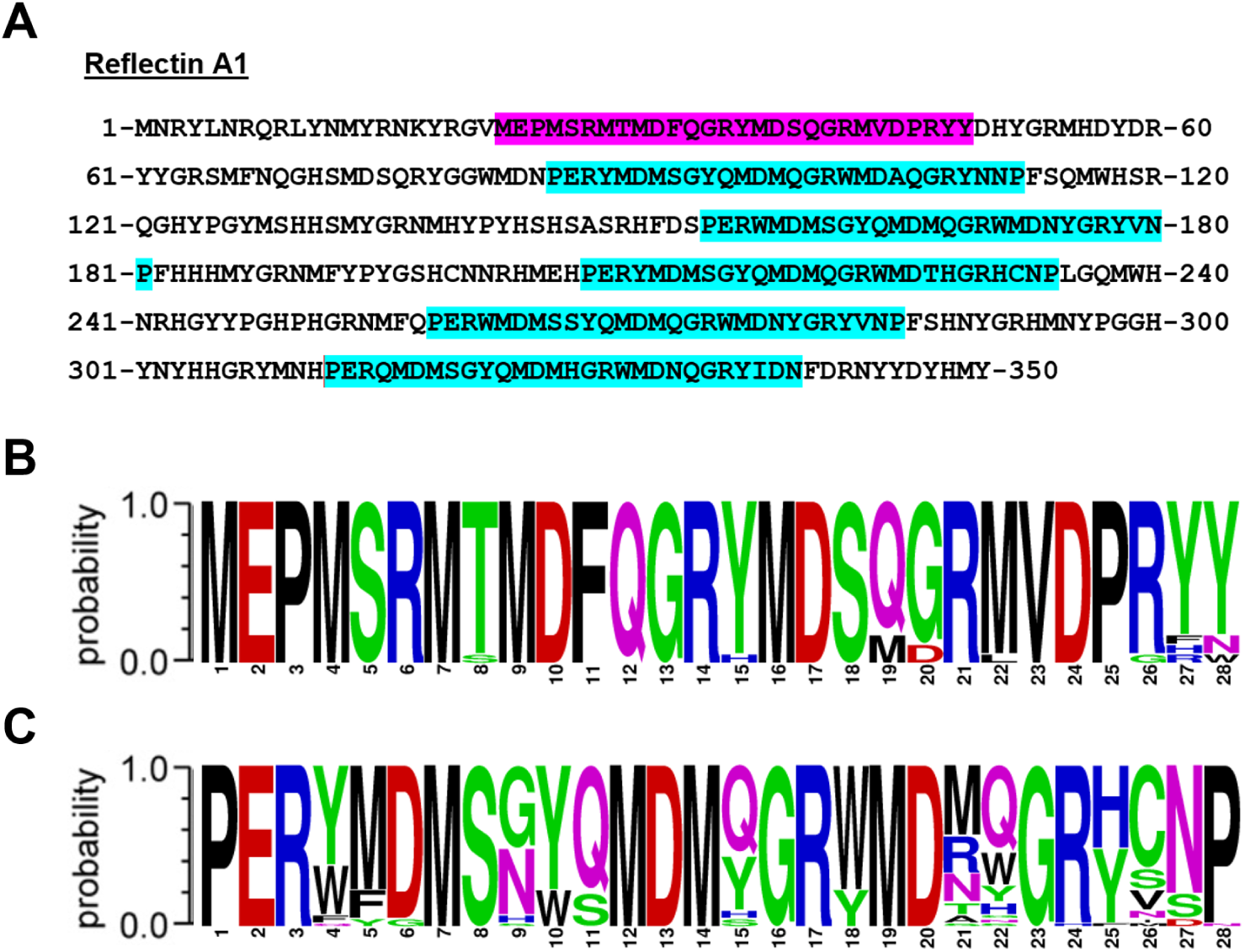
A) Amino acid sequence of reflectin A1 from *Doryteuthis opalescens* with N-terminal repeat motif (magenta) and reflectin repeat motifs (cyan). Sequence logo from alignment of the (A) N-terminal repeat motif and (B) reflectin repeat motif from 51 reflectin proteins found in *Octopus bimaculoides, Euprymna scolopes, Sepia oficinialis, Doryteuthis opalescens and Doryteuthis pealeii*.

**Figure S2.**
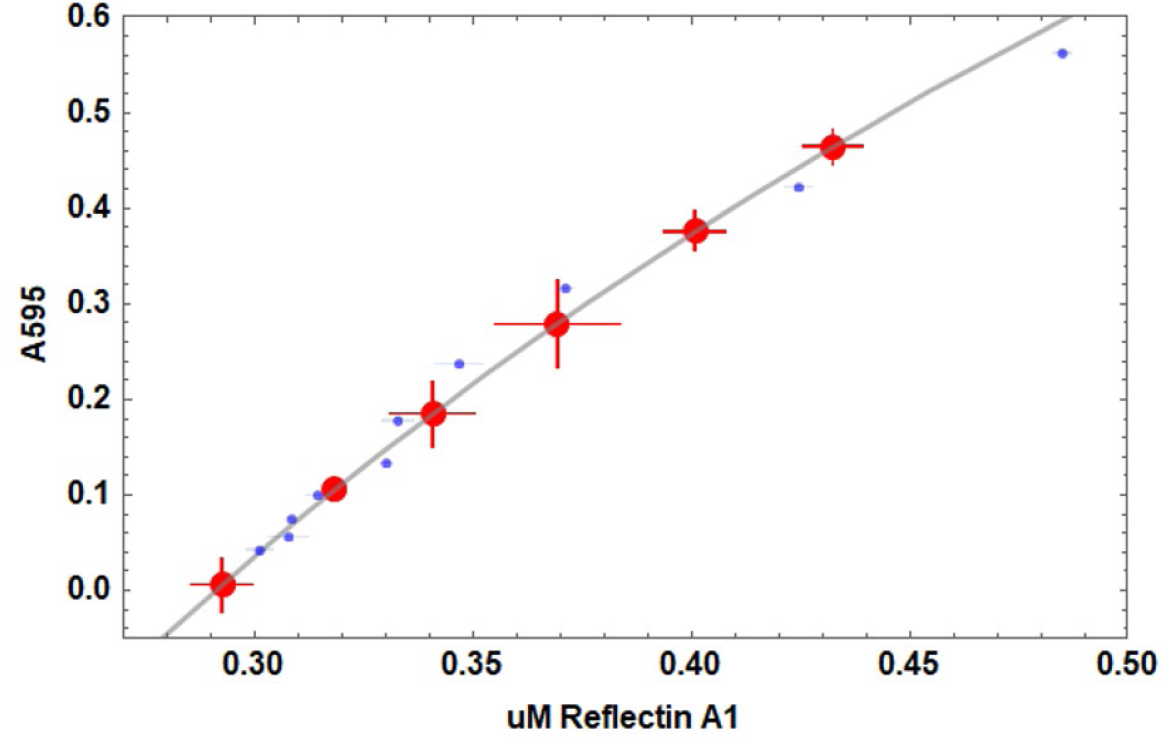
Bradford assay used to obtain values shown in Figure 1. Experimentally determined reflectin A1 concentrations using line fit (grey) to protein standards (blue). X-axis is protein concentration and Y axis is absorbance at 595 nm.

**Figure S3.**
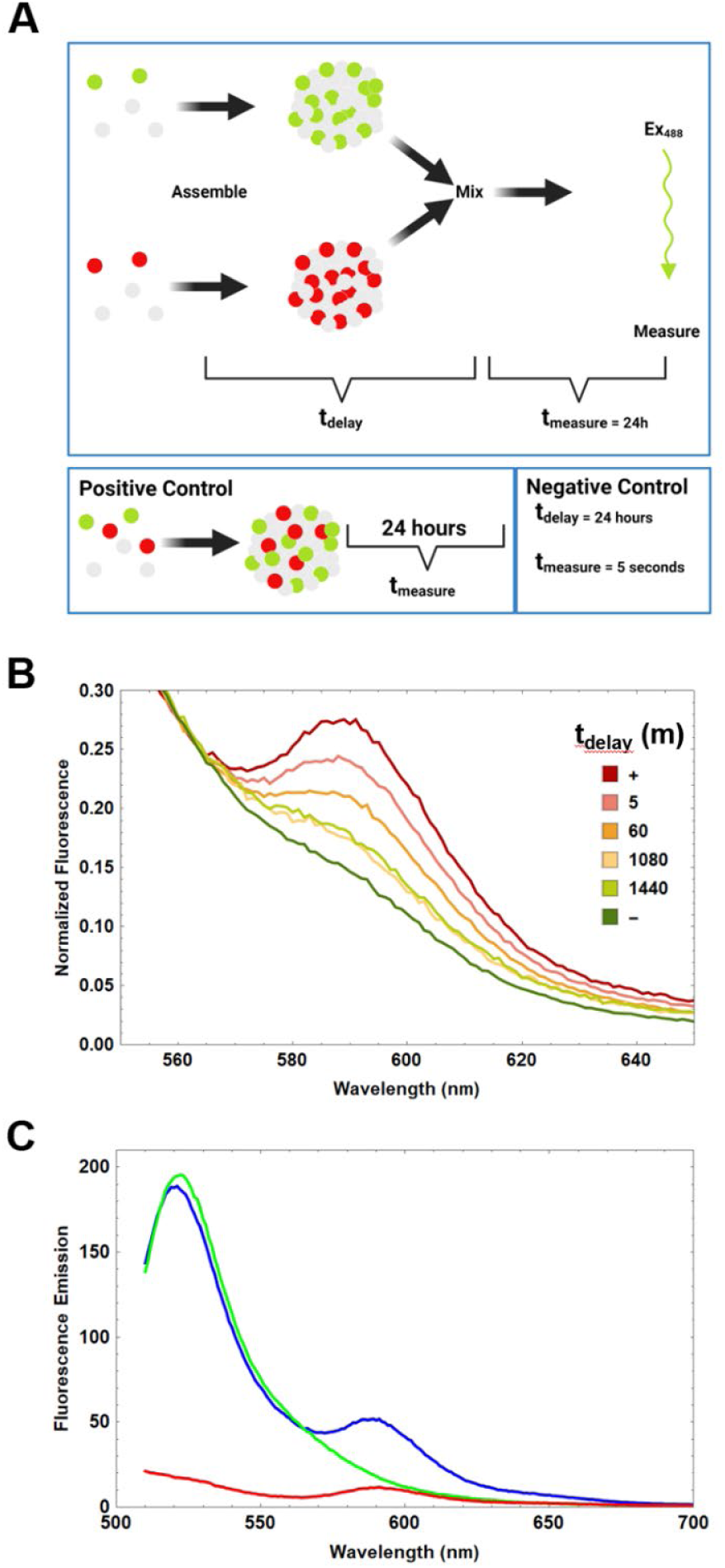
A) Design of FRET exchange experiment. Mixtures of reflectin A1 containing either 5% fluorescein-labeled single cysteine mutant C232S (A1 C199-F) or 5% rhodamine sulfate-labeled C232S (A1 C199-R) were separately then assembled by dilution into pH 7 25 mM MOPS buffer. After t_delay_ assemblies containing 5% A1 C199-F were mixed with those containing 5% A1 C199-R and incubated 24 hours at 20° C. Samples were excited with 488 nm light corresponding to the absorption spectra of donor fluorophore fluorescein and emission spectra from 510-700 nm recorded. A mixture containing both labels was driven to assembly as the positive control. For the negative control t_delay_ was 24 hours and t_measure_ was 5 seconds. B) Spectra normalized to 520 nm (fluorescein emission maximum). C) Raw fluorescence emission of reflectin A1 assemblies labeled with 5% fluorescein (green), 5% rhodamine sulfate (red) and the positive control containing both fluorescent labels (blue) excited at 488nm.

**Figure S4.**
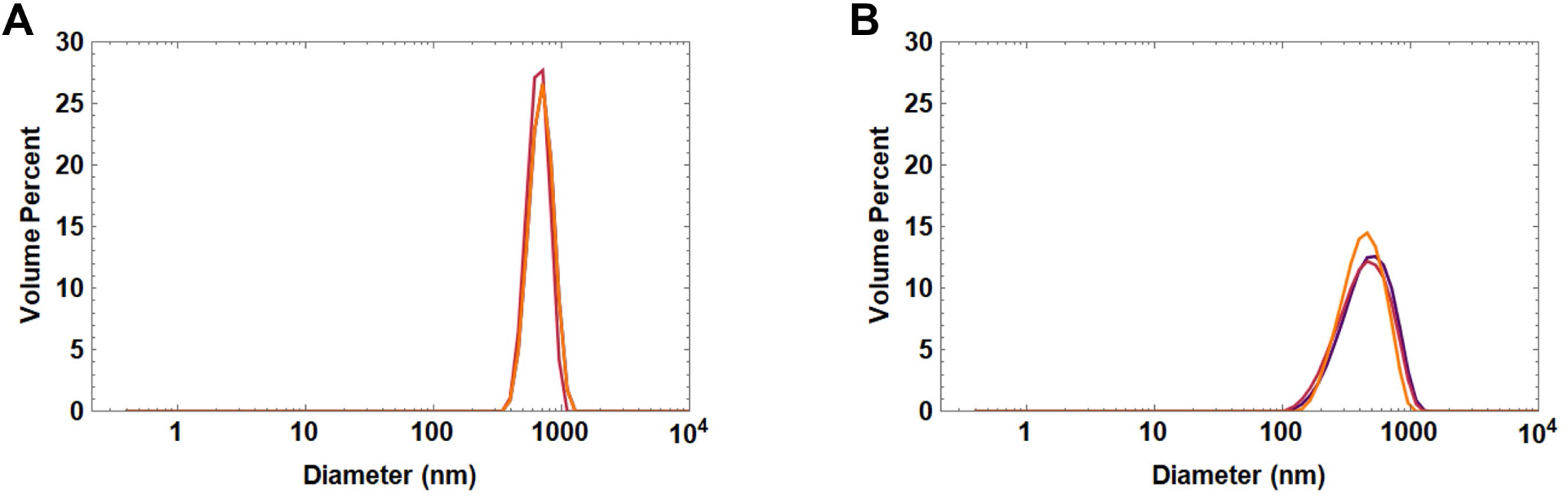
DLS size distributions of fluorescently labeled reflectin A1 C232S. A) Size distribution by volume percent of A) C199-R (rhodamine labeled reflectin A1 C232S) and B) C199-F (fluorescein labeled reflectin A1 C232S).

**Figure S5.**
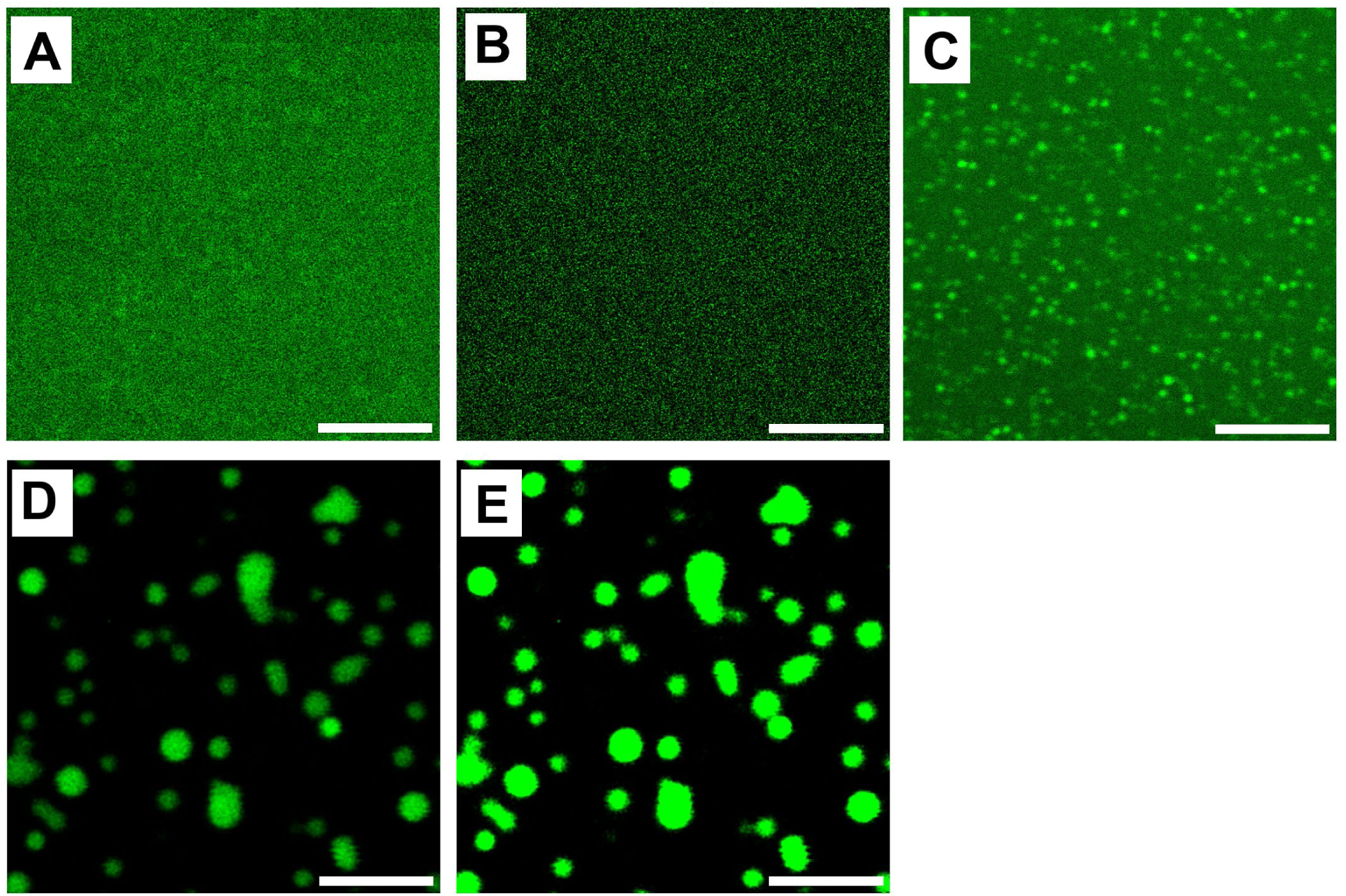
Reflectin A1 assembly sizes increase until strongly partitioned liquid droplets are formed. 100 μM reflectin A1 containing 5% A1-C199F diluted into increasing concentrations of NaCl in acetic acid buffer (pH 4 25 mM) until the liquid phase boundary is crossed. A) At 80 mM NaCl no light-resolvable structures are detected but assemblies are present as determined by DLS. B) Same as (A) but imaged using an mPEG-passivated coverslip. C) At 90 mM NaCl ca. 500 nm d. particles are present. D) At 100 mM NaCl surface-wetting liquid droplets are present. E) Same image as C but at the same exposure values as (A,B) for comparison of protein partitioning into dense phase. Scale bars = 5 µm.

**Figure S6.**
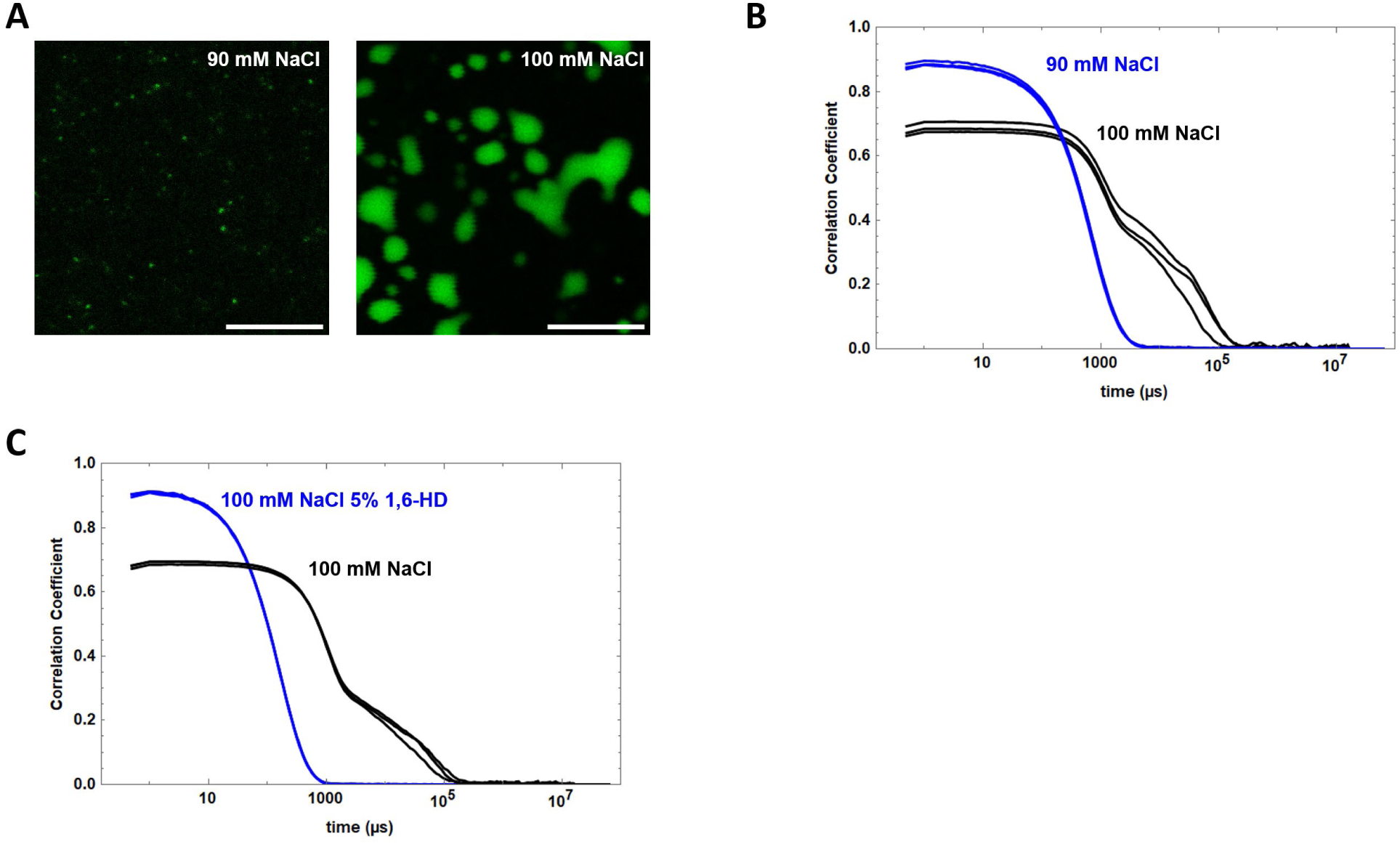
Confocal microscopy and DLS of 4 μM reflectin A1 in acetic acid buffer (pH 4, 25 mM) diluted into respective buffers. A) Confocal microscopy and (B) autocorrelation functions of reflectin A1 in 90 mM NaCl and 100 mM NaCl. C) Autocorrelation functions of reflectin A1 in 100 mM NaCl and in the same solution containing 5% 1,6-hexanediol. Autocorrelation functions are 3 successive measurement replicates of each experimental condition. Scale bars = 10 µm.

**Figure S7.**
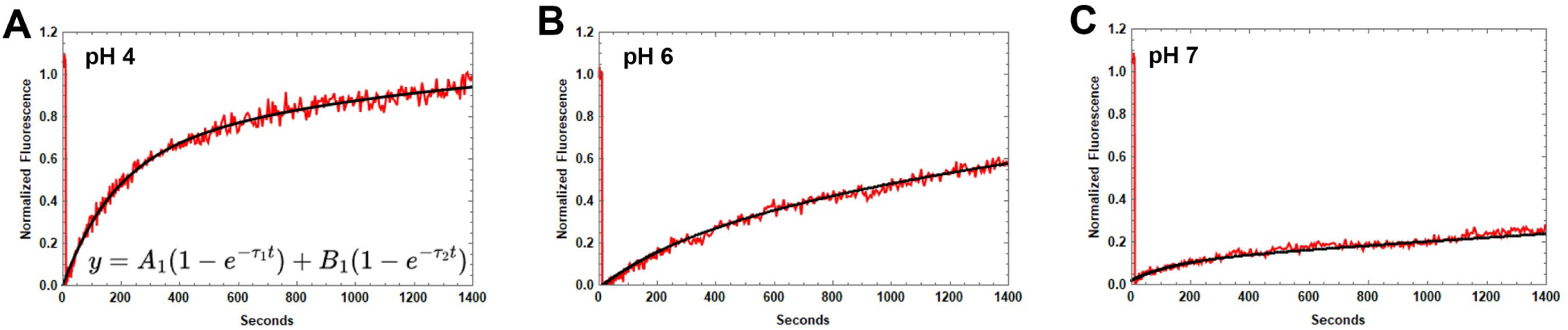
Examples of fits of individual FRAP experiments to an equation of exponential decay for (A) pH 4, (B) pH 6 and (C) pH 7.

**Figure S8.**
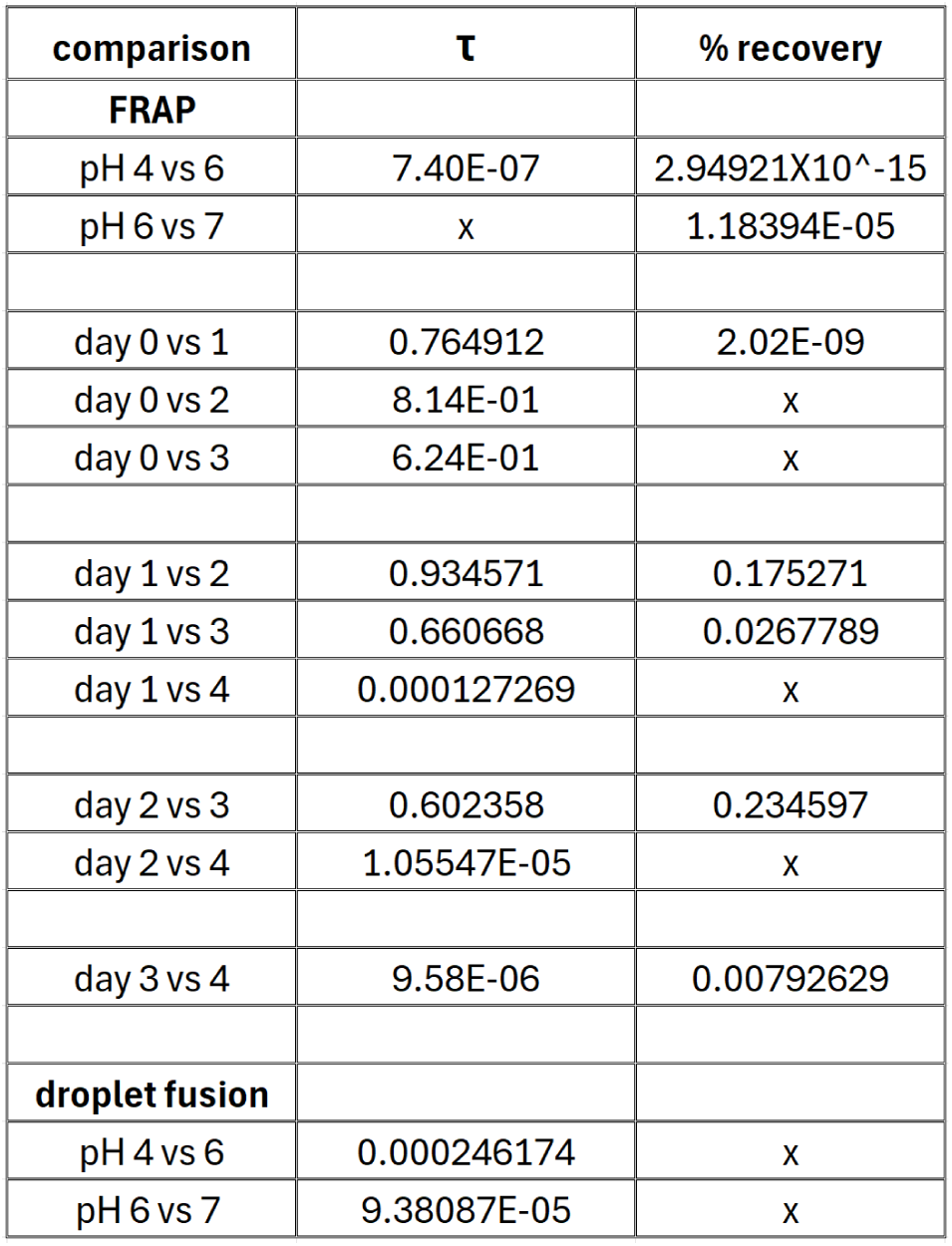
P-values obtained from statistical testing of FRAP and droplet fusion experiments. Statistical significance was determined by one-way ANOVA testing. ‘X’ denotes comparisons that were not made.

